# HTCC as a highly effective polymeric inhibitor of SARS-CoV-2 and MERS-CoV

**DOI:** 10.1101/2020.03.29.014183

**Authors:** Aleksandra Milewska, Ying Chi, Artur Szczepanski, Emilia Barreto-Duran, Kevin Liu, Dan Liu, Xiling Guo, Yiyue Ge, Jingxin Li, Lunbiao Cui, Marek Ochman, Maciej Urlik, Sylwia Rodziewicz-Motowidlo, Fengcai Zhu, Krzysztof Szczubialka, Maria Nowakowska, Krzysztof Pyrc

**Affiliations:** Virogenetics Laboratory of Virology, Malopolska Centre of Biotechnology, Jagiellonian University, Gronostajowa 7, 30-387 Krakow, Poland; Microbiology Department, Faculty of Biochemistry, Biophysics and Biotechnology, Jagiellonian University, Gronostajowa 7, 30-387 Krakow, Poland; NHC Key Lab of Enteric Pathogenic Microbiology, Jiangsu Provincial Centre for Disease Control & Prevention. 172 Jiangsu Rd., Nanjing, Jiangsu, 210009, PR China; Nanjing Techboon Institute of Clinical Medicine. #1003, Tower B of Yangtze Sci. & Tech. Innovation Centre, 211 Pubing Rd., Nanjing, Jiangsu, 211800, PR China; Department of Cardiac, Vascular and Endovascular Surgery and Transplantology, Medical University of Silesia in Katowice, Silesian Centre for Heart Diseases, Zabrze, Poland; Department of Biomedical Chemistry, Faculty of Chemistry, University of Gdansk, Wita Stwosza 63, 80-308 Gdansk, Poland; Department of Physical Chemistry, Faculty of Chemistry, Jagiellonian University, Gronostajowa 2, 30-387, Krakow, Poland; Centre for Global Health, Nanjing Medical University. 18 Tianyuan Rd. E., Nanjing, Jiangsu, 210009, PR China

## Abstract

The beginning of 2020 brought us information about the novel coronavirus emerging in China. Rapid research resulted in the characterization of the pathogen, which appeared to be a member of the SARS-like cluster, commonly seen in bats. Despite the global and local efforts, the virus escaped the healthcare measures and rapidly spread in China and later globally, officially causing a pandemic and global crisis in March 2020. At present, different scenarios are being written to contain the virus, but the development of novel anticoronavirals for all highly pathogenic coronaviruses remains the major challenge. Here, we describe the antiviral activity of previously developed by us HTCC compound (N-(2-hydroxypropyl)-3-trimethylammonium chitosan chloride), which may be used as potential inhibitor of currently circulating highly pathogenic coronaviruses – SARS-CoV-2 and MERS-CoV.

## INTRODUCTION

Coronaviruses mainly cause respiratory and enteric diseases in humans, other mammals, and birds. However, some species can cause more severe conditions such as hepatitis, peritonitis, or neurological disease. Seven coronaviruses infect humans, four of which (human coronavirus [HCoV]-229E, HCoV-NL63, HCoV-OC43, and HCoV-HKU1) cause relatively mild upper and lower respiratory tract disease. Other three zoonotic coronaviruses - the severe acute respiratory syndrome coronaviruses (SARS-CoV and SARS-CoV-2) and the Middle East respiratory syndrome coronavirus (MERS-CoV) are associated with severe, life-threatening respiratory infections and multiorgan failure ^1–7^.

SARS-CoV-2 emerged in the Hubei province of China by the end of 2019 and caused an epidemic that was partially contained in China by March 2020. However, the virus rapidly spread globally and caused the pandemic ^8^. SARS-CoV-2 is a betacoronavirus and belongs to a large cluster of SARS-like viruses in bats, classified in *Sarbecovirus* subgenus. While bats are considered to be the original reservoir, it is believed that there is an intermediate host, and pangolins were suggested as such ^9^. The virus is associated with a respiratory illness that, in a proportion of cases, is severe. The mortality rate varies between locations, but at present, is estimated to reach 3-4% globally. The virus infects primarily ciliated cells and type II pneumocytes in human airways, hijacking the angiotensin-converting enzyme 2 (ACE2) to enter the cell, similarly as SARS-CoV and HCoV-NL63.

MERS-CoV is related to SARS-CoV-2, but together with some bat viruses forms a separate *Merbecovirus* subgenus. Bats are believed to serve as an original reservoir also in this case ^10^, but camels were identified as the intermediate host ^11^. The virus never fully crossed the species border, as the human-to-human transmission is limited, and almost all the cases are associated with animal-to-human transmission. The entry receptor for MERS-CoV is the dipeptidyl peptidase 4 (DPP4) ^12,13^. In humans, MERS-CoV causes a respiratory illness with severity varying from asymptomatic to potentially fatal acute respiratory distress ^14–16^. To date, MERS-CoV infection was confirmed in 27 countries, with over 2,000 cases and a mortality rate of ~35%.

Currently, there are no vaccines or drugs with proven efficacy to treat coronavirus infection, and treatment is limited to supportive care. However, a range of therapeutics have been experimentally used in clinic to treat SARS-CoV-2 and MERS-CoV-infected patients, and their use is based on the knowledge obtained in previous years. The most promising drug candidates include broad-spectrum polymerase inhibitors (remdesivir) ^17^ and some re-purposed drugs (e.g., HIV-1 protease inhibitors). However, until today none of these has proven effective in randomized controlled trials. On the other hand, the antiviral potential of several small molecules have been demonstrated in cell lines *in vitro*, but their effectiveness *in vivo* have not been confirmed ^18,19^. For the ones that reached animal models, some promising drug candidates have been shown to exacerbate the disease (ribavirin, mycophenolic acid) ^20^.

Drug development for SARS-CoV-2 and MERS-CoV is of great importance, but considering the diversity of coronaviruses and the proven propensity to cross the species barrier, the SARS-CoV-2 epidemic is probably not the last one. Warnings about the possibility of another SARS-CoV epidemic have appeared in the scientific literature for a long time ^21^. Consequently, broad-spectrum antivirals are essential in long-term perspective.

Previously, we have demonstrated an antiviral potential of the HTCC polymer (N-(2-hydroxypropyl)-3-trimethylammonium chitosan chloride), which efficiently hampered infection of all low pathogenic human coronaviruses *in vitro* and *ex vivo* ^22^ and several animal coronaviruses (*unpublished data*). Furthermore, using several functional and structural assays, we dissected the mechanism of the HTCC antiviral activity. We showed that the polymer interacts with the coronaviral Spike (S) protein and blocks its interaction with the cellular receptor ^22–24^. Here, we analyzed the HTCC activity against SARS-CoV-2 and MERS-CoV *in vitro* using permissive cell lines and *ex vivo*, using a model of human airway epithelium (HAE). The study showed that the replication of both viruses was efficiently hampered. Overall, our data show that HTCC polymers are potent broad-spectrum anticoronavirals and may be very promising drug candidates for SARS-CoV-2 and MERS-CoV.

## RESULTS AND DISCUSSION

### HTCCs hamper MERS-CoV and SARS-CoV-2 replication in cell lines

Previously, we showed that HTCC with different degrees of substitution (DSs) is a potent inhibitor of all four low pathogenic HCoVs ^22^. DSs are expressed as the fraction of NH_2_ groups of gluocosamine units in chitosan substituted with glycidyltrimethylammonium chloride (GTMAC). The DS of the studied HTCC polymers varied between 57% and 77%; thus, the polymers were named HTCC-57, HTCC-62, HTCC-63, HTCC-65, and HTCC-77. The synthesis and characterization of polymers are described elsewhere ^22,23^. The analysis showed that HTCC-63 demonstrated the most significant inhibitory effect on HCoV-NL63, HCoV-OC43, and HCoV-HKU1. On the other hand, HTCC-62 and HTCC-77 proved to be effective inhibitors of HCoV-229E infection. Further, HTCC-65 effectively inhibited the replication of HCoV-NL63 and HCoV-OC43, while HTCC-62 showed a potent antiviral effect on HCoV-HKU1 infection.

The study on the MERS-CoV using the Vero cells revealed that all HTCC variants inhibit virus replication to a similar extent (~100-1000-time decrease in viral yields), at non-toxic concentration (**Figure 1A, C**). The inhibition of the SARS-CoV-2 infection in Vero E6 cells was even more pronounced, and all HTCC variants inhibited virus replication by ~10,000 times at non-toxic concentration (**Figure 1B, C**). In this case, the HTCC-63 was arbitrarily selected for further studies on MERS-CoV, while HTCC-77 was selected for SARS-CoV-2.

**Figure 1.**
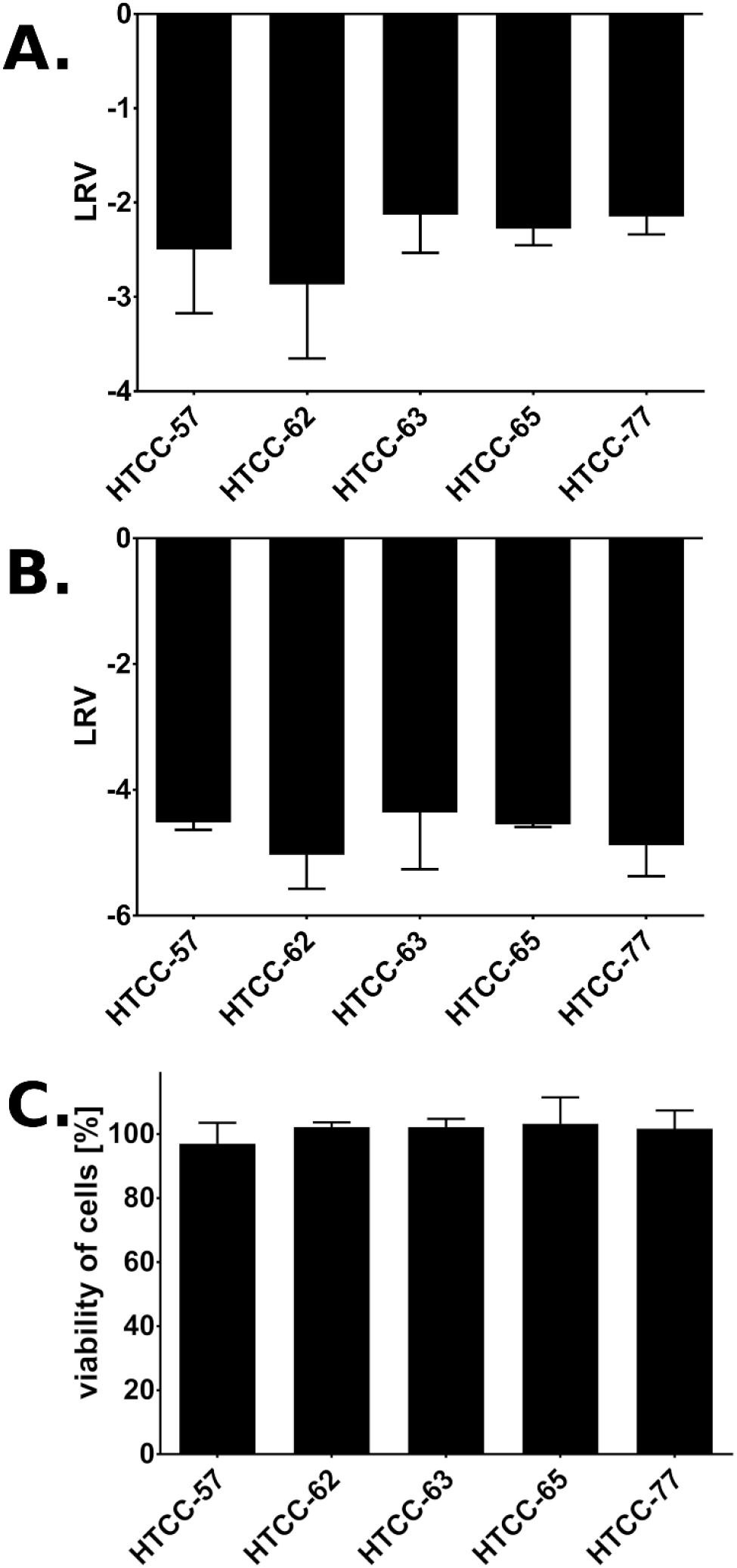
*In vitro* inhibition of MERS-CoV and SARS-CoV-2 by HTCC at non toxic concentration. Vero cells were infected with MERS-CoV (**A**), and Vero E6 cells were infected with SARS-CoV-2 (**B**). Briefly, cultures were inoculated with a given virus in the presence of HTCC (100 μg/ml) or control PBS. Replication of viruses was evaluated at 48 h post-inoculation using RT-qPCR. The data are presented as Log Removal Value (LRV) compared to the untreated sample. The assay was performed in triplicate, and average values with standard errors are shown. (**C**) Cytotoxicity of HTCCs with DS ranging from 57% to 77% *in vitro* at 100 μg/ml. Cell viability was assessed with XTT assay. Data on the y-axis represent the percentage of values obtained for the untreated reference samples. All assays were performed in triplicate, and average values with standard errors are presented. The differences in cytotoxicity of HTCCs were not statistically significant.

Next, the dose-dependence was tested for the HTCCs. The inhibitory activity of selected polymers was verified for three different concentrations, and obtained data are shown in **Figure 2**.

**Figure 2.**
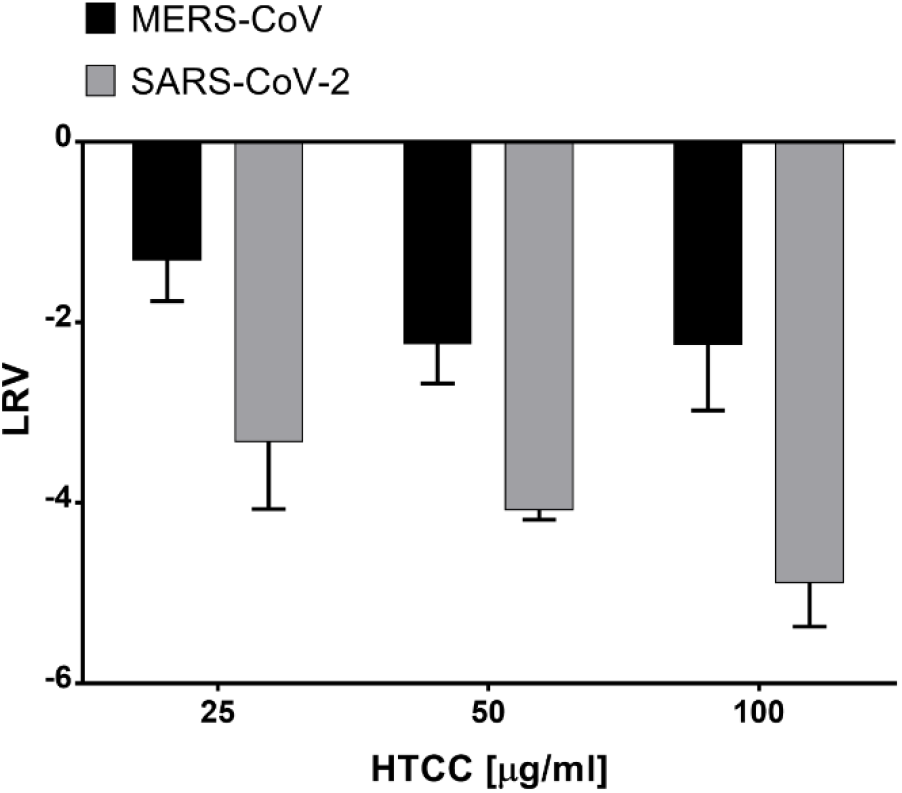
Dose-dependent inhibition of MERS-CoV and SARS-CoV-2 replication. Vero (MERS-CoV) or Vero E6 cells (SARS-CoV-2) were inoculated with a given virus in the presence of different concentrations of HTCC. Replication of viruses was evaluated at 48 h post-inoculation using RT-qPCR. The data are presented as Log Removal Value (LRV) compared to the untreated sample. The assay was performed in triplicate, and average values with standard errors are displayed.

Based on the data obtained, the basic parameters were calculated and are presented in **Table 1**.

**Table 1.**
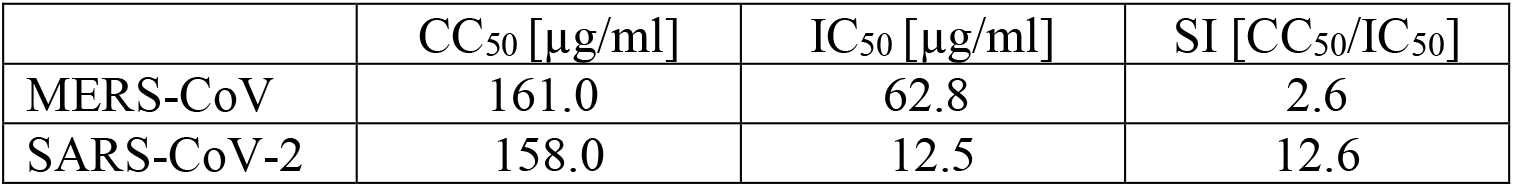
50% Cytotoxic concentration (CC_50_), 50% inhibitory concentration (IC_50_), and the selectivity index (SI) of two most effective HTCCs: HTCC-63 (for MERS-CoV) and HTCC-77 (for SARS-CoV-2).

The parameters observed for the SARS-CoV-2 appear to be favorable with SI above 12. It is also worth to note that HTCC was previously administered by inhalation in rats, and no adverse reactions were observed ^25^. In that study, HTCC was used as a carrier for the active substance, and as such, was reported to be promising for local sustained inhalation therapy of pulmonary diseases.

### HTCCs hamper MERS-CoV and SARS-CoV-2 replication in human airway epithelium

While the Vero cells constitute a convenient model for antiviral research, it is of utmost importance to verify whether the results obtained are not biased due to the artificial system used. This is especially important for compounds, which activity is based on electrostatic interaction. To verify whether the natural microenvironment, which is rich in sugars and charged molecules, does not abrogate the effectiveness of the inhibitors, we employed HAE cultures that mirror the fully differentiated layer linng the conductive airways, as well as the site of coronavirus replication. Briefly, fully differentiated HAE cultures were infected with a given virus ^26^ in the presence of previously selected HTCCs (200 μg/ml) or control PBS. Following inoculation, apical lavage samples were collected daily, and replication kinetics for each virus was investigated. The analysis revealed that the polymer efficiently hampered SARS-CoV-2 and MERS-CoV also in this model. For MERS-CoV, the inhibitory effect was the most evident at 72 h p.i., while for SARS-CoV-2 the most substantial decline of virus progeny was observed at 24 h p.i. (**Figure 3**). Whether such kinetics will be reflected *in vivo* it is to be investigated.

**Figure 3.**
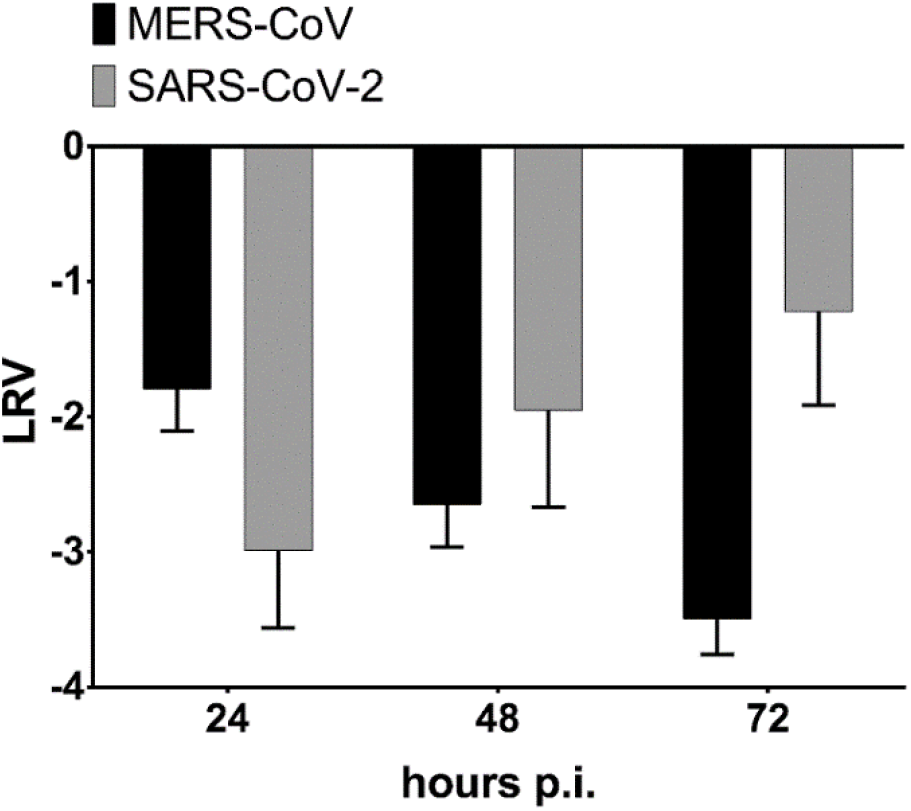
*Ex vivo* inhibition of MERS-CoV and SARS-CoV-2 by HTCC in human airway epithelium cultures. HAE cultures were exposed to MERS-CoV or SARS-CoV-2 in the presence of HTCC-63 (for MERS-CoV) or HTCC-77 (for SARS-CoV-2) at 200 μg/ml or control PBS. To analyze virus replication kinetics, each day post infection, 100 μl of 1 × PBS was applied to the apical surface of HAE cultures and collected after 10 min of incubation at 37°C. Replication of viruses was evaluated using quantitative RT-qPCR. The data are presented as Log Removal Value (LRV) compared to the untreated sample. The assay was performed in triplicate, and average values with standard errors are shown.

### HTCCs inhibits MERS-CoV and SARS-CoV-2 entry into susceptible cells

Our previous research showed that the HTCC-mediated inhibition of coronaviral replication results from the electrostatic interaction between the polymer and the Spike protein of coronaviruses. We hypothesize that the selectivity of the inhibitors yields from the fitting charge distributions on the polymer and on the S proteins on the viral surface. While the interaction of a single charged moiety is relatively weak, the concatemeric nature of the virus and the polymer stabilizes the binding. Such structure-based interaction may be an interesting entry point for further fine-tuning of the polymeric inhibitors of viral replication.

To ensure that the observed effect was a result of coronavirus entry inhibition by HTCC, two experiments were performed. First, HAE cultures were inoculated with MERS-CoV in the presence of HTCC-63 (200 μg/ml) or control PBS and incubated for 2 h at 37°C. Next, cells were fixed, immunostained for MERS-CoV N protein and actin. Virus entry was analyzed with confocal microscopy. To visualize the effect, the signal attributed to intracellular MERS-CoV was quantified, and the results show that the internalization of MERS-CoV was drastically decreased (**Figure 4A**).

**Figure 4.**
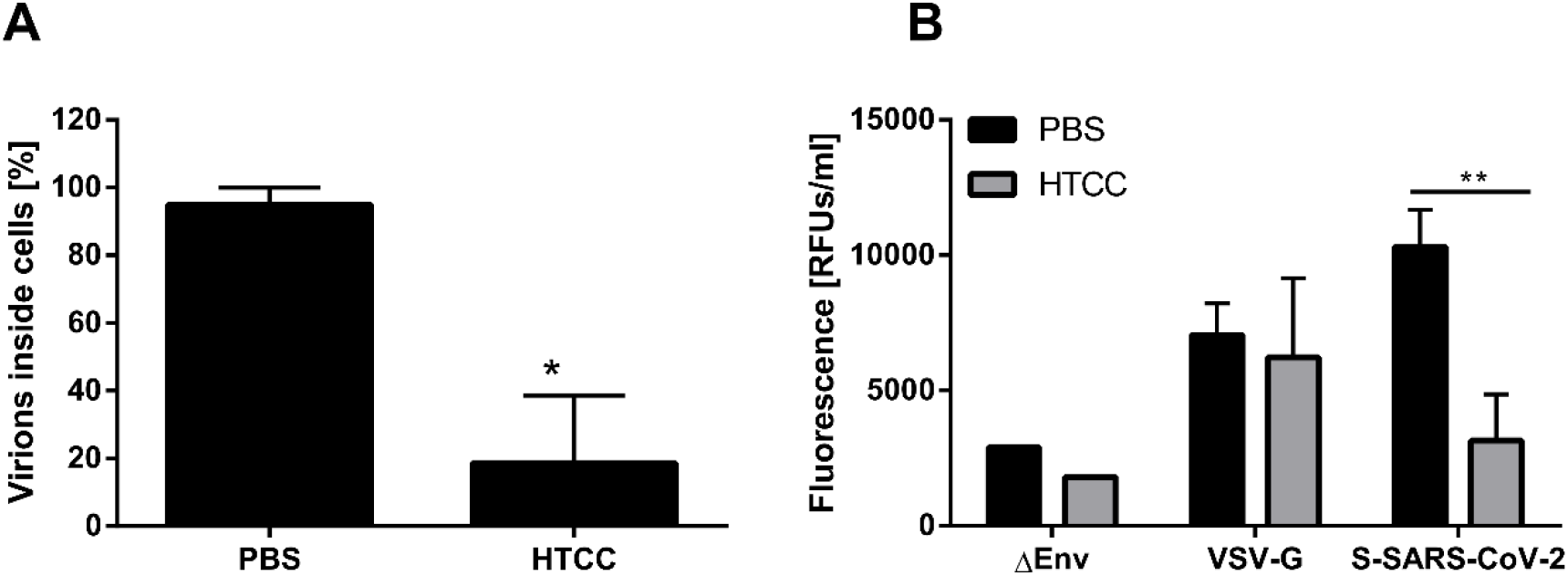
Coronavirus internalization into susceptible cells is hampered by HTCC. **(A)** Pre-cooled HAE cultures were incubated with ice-cold MERS-CoV suspension in the presence or absence of HTCC-63 (200 μg/ml) for 2 h at 37°C. Next, cells were fixed in PFA and immunostained for MERS-CoV N protein and actin. Virus entry was analyzed with confocal microscopy. The data shown are representative of three independent experiments, each performed in triplicate. * P < 0.05. (**B**) A549 cells overexpressing ACE2 were incubated with lentiviral particles bearing GFP reporter gene, pseudotyped with SARS-CoV-2 Spike (S-SARS-CoV-2), VSV control G protein (VSV-G) or particles without an envelope protein (∆Env) in the presence of HTCC-77 or control PBS. After 2 h at 37°C cells were washed with PBS and overlaid with fresh medium. Following 72 h incubation, GFP signal was measured using fluorometer and pseudovirus entry is presented as Relative Fluorescence Units per ml. The assay was performed in triplicate, and average values with standard errors are presented. ** P < 0.005.

Due to the limited availability of tools for the SARS-CoV-2 we were not able to replicate the experiment for this virus. Here, we employed a surrogate system based on lentiviral vectors pseudotyped with full-length Spike protein of SARS-CoV-2. A549 cells overexpressing the ACE2 protein were incubated with pseudovirions harboring SARS-CoV-2 Spike or control VSV-G protein in the presence of HTCC-77 (100 μg/ml) or control PBS for 2 h at 37°C. After 72 h p.i. cells were lysed, and pseudovirus entry was quantified by measurement of the reporter GFP protein. The analysis showed a significant reduction in SARS-CoV-2 Spike pseudoviruses internalization in the presence of the polymer, while no inhibition was observed for the control VSV-G (**Figure 4B**).

Next, to verify whether the mechanism of action for the highly pathogenic betacoronaviruses is similar to that observed for alphacoronaviruses ^22^, and is based on locking the interaction between the virus and the entry receptor, we analyzed MERS-CoV co-localization with its entry receptor, DPP4 in the presence or absence of the HTCC. For this, human cell line Huh7 was inoculated with the virus or mock in the presence of HTCC-63 (100 μg/ml) or control PBS and incubated for 2 h at 4°C. Subsequently, cells were fixed and immunostained for the DPP4 and MERS-CoV N using specific antibodies. Virus co-localization with its receptor was examined using confocal microscopy. Obtained results demonstrated that in the control samples, virions co-localize with the DPP4 protein, while in the presence of the polymer, this interaction is blocked (**Figure 5**).

**Figure 5.**
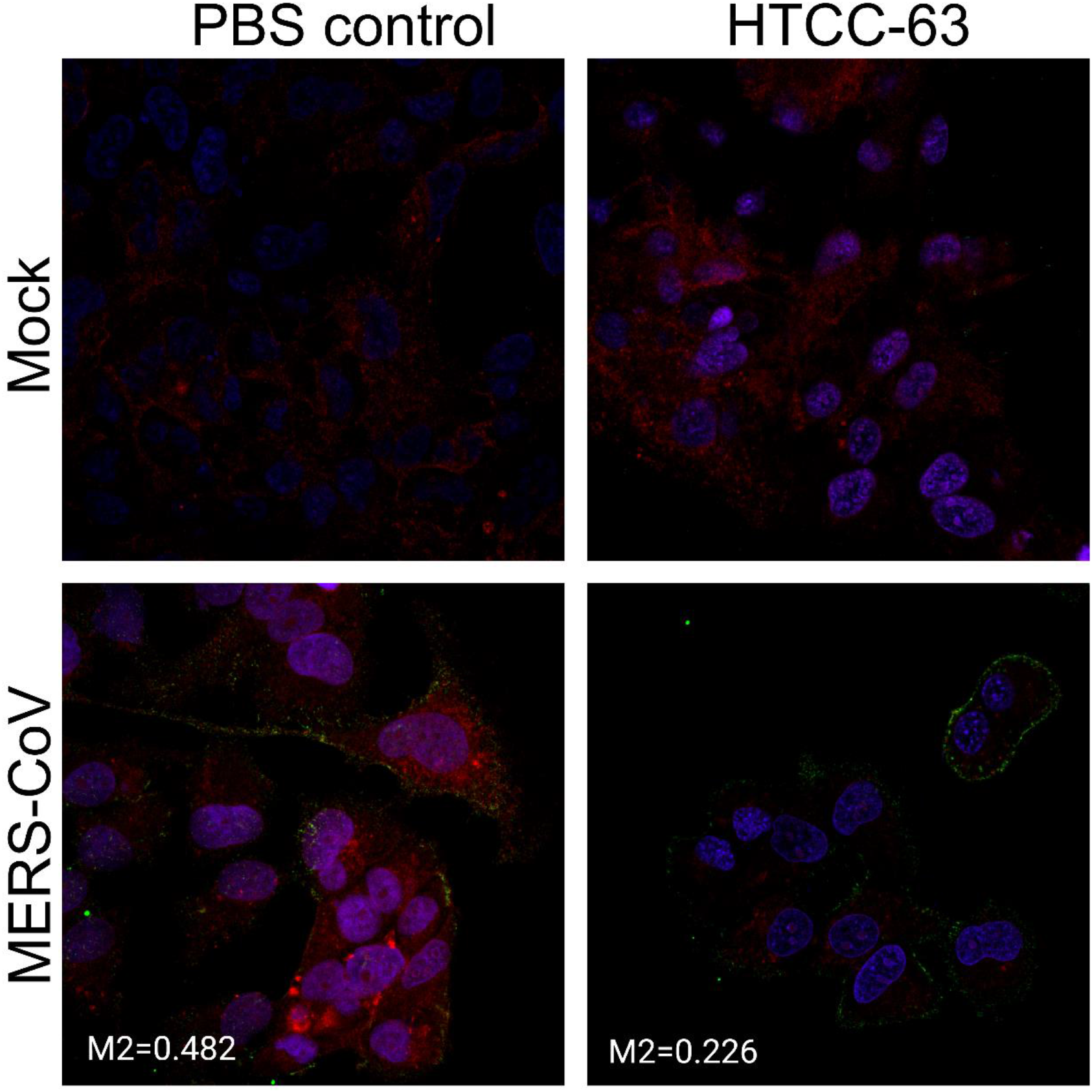
HTCC blocks interaction between the virus and its entry receptor. Pre-cooled Huh7 cells were incubated for 3 h at 4°C with ice-cold MERS-CoV or mock in the presence or absence of HTCC-63 (100 μg/ml). Next, cells were fixed with PFA and immunostained for MERS-CoV-N (green), DPP4 (red) and nuclear DNA (blue). MERS-CoV interaction with the DPP4 protein was analyzed with confocal microscopy. Co-localisation of DPP4 with MERS-CoV-N was determined by confocal microscopy and is presented as Manders’ M2 coefficient. The decrease in colocalization was statistically significant (P < 0.0005). Each image represents maximum projection of axial planes. Representative images are shown.

Taking together, we show here that the previously developed and described polymeric HTCC anticoronaviral compounds based on chitosan are able to efficiently inhibit infection with emerging coronaviruses. We believe that the HTCC can be fine-tuned to target any coronavirus, and this interaction is specific to viruses that belong to the *Coronaviridae* family. One may speculate that the inhibition results from the concatemeric nature of the virus surface and the fact that the polymer with appropriate charge distribution can interact with multiple sites on this surface. While the interaction of the monomer is relatively weak, and no inhibition is observable for monomers, the sum of interactions stabilizes the binding and specific inhibition is observed. Considering that the extended chain length for the HTCC used is ~700 nm this scenario seems realistic ^27^. If that would be true, HTCC would constitute a first structure-specific inhibitor of viral replication. The major disadvantage of the HTCC is that, at present, it is not registered for use in humans. However, previous experience with HTCC in different laboratories shows that it may be delivered topically to the lungs, it is not associated with toxicity, and it does not worsen the lung function ^25^. We believe that HTCC is a promising drug candidate that should be further studied, as it provides a ready-to-use solution for SARS-CoV-2 and future emerging coronaviruses.

## MATERIALS AND METHODS

### The active compound

The HTCC was prepared in the same manner as previously described ^22,23,28^.

### Plasmid constructs

The codon-optimised full-length SARS-CoV-2 S gene was designed and purchased from GeneArt (Thermo Fisher Scientific, Poland). The gene was cloned into pCAGGS vector sequence verified that was a gift from Xingchuan Huang. psPAX (Addgene plasmid # 12260) and pMD2G (Addgene plasmid # 12259) was a gift from Didier Trono. Lego-G2 vector (Addgene plasmid #25917) was a gift from Boris Fehs.

### Cell culture

Vero and Vero E6 (*Cercopithecus aethiops*; kidney epithelial; ATCC: CCL-81 and CRL-1586), Huh7 (*Homo sapiens*; hepatocellular carcinoma; ECACC: 01042712) and A549 cells with ACE2 overexpression (A549/ACE2)^29^ were cultured in Dulbecco’s MEM (Thermo Fisher Scientific, Poland) supplemented with 3% fetal bovine serum (heat-inactivated; Thermo Fisher Scientific, Poland) and antibiotics: penicillin (100 U/ml), streptomycin (100 μg/ml), and ciprofloxacin (5 μg/ml). Cells were maintained at 37°C under 5% CO_2_.

### Human airway epithelium (HAE) cultures

Human airway epithelial cells were isolated from conductive airways resected from transplant patients. The study was approved by the Bioethical Committee of the Medical University of Silesia in Katowice, Poland (approval no: KNW/0022/KB1/17/10 dated 16.02.2010). Written consent was obtained from all patients. Cells were dislodged by protease treatment, and later mechanically detached from the connective tissue. Further, cells were trypsinized and transferred onto permeable Transwell insert supports (ϕ = 6.5 mm). Cell differentiation was stimulated by the media additives and removal of media from the apical side after the cells reached confluence. Cells were cultured for 4-6 weeks to form well-differentiated, pseudostratified mucociliary epithelium. All experiments were performed in accordance with relevant guidelines and regulations.

### Cell viability assay

HAE cultures were prepared as described above. Cell viability assay was performed by using the XTT Cell Viability Assay (Biological Industries, Israel) according to the manufacturer’s instructions. On the day of the assay, 100 μl of the 1 × PBS with the 50 μl of the activated XTT solution was added to each well/culture insert. Following 2 h incubation at 37°C, the solution was transferred onto a 96-well plate, and the signal was measured at λ = 490 nm using the colorimeter (Spectra MAX 250, Molecular Devices). The obtained results were further normalized to the control sample, where cell viability was set to 100%.

### Virus preparation and titration

MERS-CoV stock (isolate England 1, 1409231v, National Collection of Pathogenic Viruses, Public Health England, United Kingdom) was generated by infecting monolayers of Vero cells. SARS-CoV-2 stock (isolate 026V-03883; kindly granted by Christian Drosten, Charité – Universitätsmedizin Berlin, Germany by the European Virus Archive - Global (EVAg); https://www.european-virus-archive.com/) was generated by infecting monolayers of Vero E6 cells. The virus-containing liquid was collected at day 3 post-infection (p.i.), aliquoted and stored at −80°C. Control Vero or Vero E6 cell lysate from mock-infected cells was prepared in the same manner. Virus yield was assessed by titration on fully confluent Vero or Vero E6 cells in 96-well plates, according to the method of Reed and Muench. Plates were incubated at 37°C for 3 days and the cytopathic effect (CPE) was scored by observation under an inverted microscope.

### Virus infection

In *in vitro* experiments, fully confluent Vero, Vero E6, or Huh7 cells in 96-well plates (TPP) were exposed to MERS-CoV, SARS-CoV-2 or mock at a TCID_50_ of 400 per ml in the presence of tested polymer or control medium. Following a 2 h incubation at 37°C, unbound virions were removed by washing with 100 μl of 1 × PBS and fresh medium containing dissolved respective polymer was added to each well. Samples of cell culture supernatant were collected at day 3 p.i. and analyzed using RT-qPCR.

For the *ex vivo* study, fully differentiated human airway epithelium (HAE) cultures were exposed to the tested polymer or control PBS for 30 min at 37°C, following inoculation with MERS-CoV or SARS-CoV-2 at a TCID_50_ of 400 per ml in the presence of the polymer or control PBS. Following 2 h incubation at 37°C, unbound virions were removed by washing with 200 μl of 1 × PBS and HAE cultures were maintained at an air—liquid interphase for the rest of the experiment. To analyze virus replication kinetics, each day p.i., 100 μl of 1 × PBS was applied at the apical surface of HAE and collected following the 10 min incubation at 32°C. All samples were stored at −80°C and analyzed using RT-qPCR.

### Isolation of nucleic acids and reverse transcription (RT)

Viral DNA/RNA Kit (A&A Biotechnology, Poland) was used for nucleic acid isolation from cell culture supernatants, according to the manufacturer’s instructions. cDNA samples were prepared with a High Capacity cDNA Reverse Transcription Kit (Thermo Fisher Scientific, Poland), according to the manufacturer’s instructions.

### Quantitative PCR (qPCR)

Viral RNA yield was assessed using real-time PCR (7500 Fast Real-Time PCR; Life Technologies, Poland). cDNA was amplified in a reaction mixture containing 1 × qPCR Master Mix (A&A Biotechnology, Poland), in the presence of probe (100 nM) and primers (450 nM each).

**Table 1.**
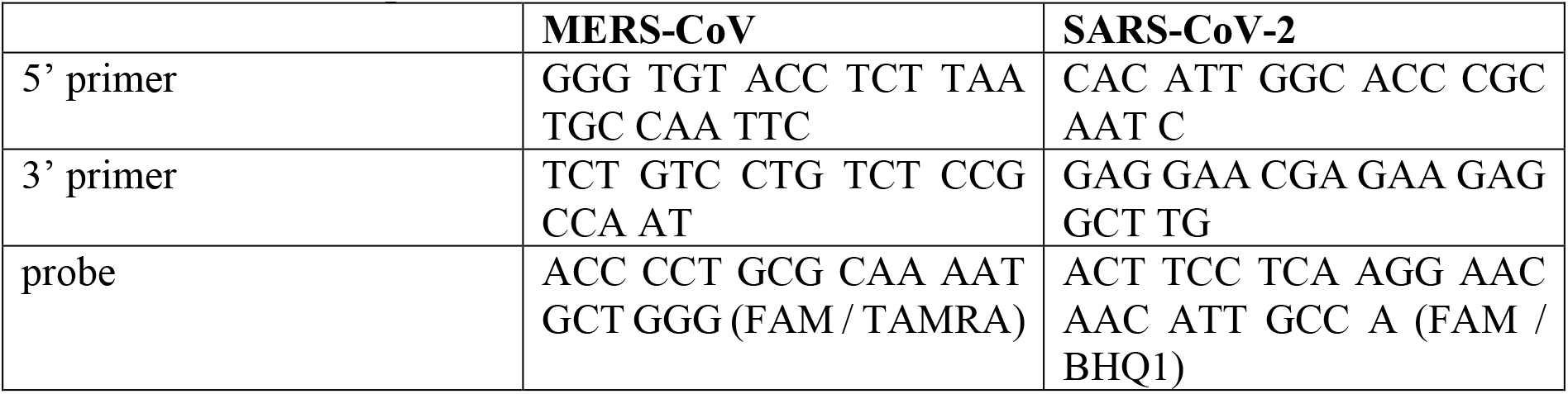
Primers and probes.

The reaction was carried out according to the scheme: 2 min at 50°C and 10 min at 92°C, followed by 40 cycles of 15 s at 92°C and 1 min at 60°C. In order to assess the copy number for N gene, DNA standards were prepared, as described before ^26^.

### Immunostaining and confocal imaging

Fixed cells were permeabilized with 0.1% Triton X-100 in 1 × PBS and incubated overnight at 4°C in 1× PBS supplemented with 5% bovine serum albumin (BSA) and 0.5% Tween 20. To visualize MERS-CoV particles, cells were incubated for 2 h at room temperature with mouse anti-MERS-CoV N IgGs (1:000 dilution, Sino Biological, China), followed by 1 h of incubation with Alexa Fluor 488-labeled goat anti-mouse IgG (2.5 µg/ml; Thermo Fisher Scientific, Poland). Actin filaments was stained using phalloidin coupled with Alexa Fluor 633 (0.2 U/ml; Thermo Fisher Scientific, Poland). Nuclear DNA was stained with DAPI (4’,6’-diamidino-2-phenylindole) (0.1 µg/ml; Sigma-Aldrich, Poland). Immunostained cultures were mounted on glass slides in ProLong Gold antifade medium (Thermo Fisher Scientific, Poland). Fluorescent images were acquired under a Leica TCS SP5 II confocal microscope (Leica Microsystems GmbH, Mannheim, Germany) and a Zeiss LSM 710 confocal microscope (Carl Zeiss Microscopy GmbH). Images were acquired using Leica Application Suite Advanced Fluorescence LAS AF v. 2.2.1 (Leica Microsystems CMS GmbH) or ZEN 2012 SP1 software (Carl Zeiss Microscopy GmbH) deconvolved with Huygens Essential package version 4.4 (Scientific Volume Imaging B.V., The Netherlands) and processed using ImageJ 1.47v (National Institutes of Health, Bethesda, MD, USA). At the time of the study, no antibodies specific to SARS-CoV-2 were available to us.

### Pseudovirus production and transduction

293T cells were seeded on 10 cm^2^ dishes, cultured for 24 h at 37°C with 5% CO_2_ and transfected using polyethyleneimine (Sigma-Aldrich, Poland) with the lentiviral packaging plasmid (psPAX), the VSV-G envelope plasmid (pMD2G) or SARS-CoV-2 S glycoprotein (pCAGGS-SARS-CoV-2-S) and third plasmid encoding GFP protein (Lego-G2). Cells were further cultured for 72 h at 37°C with 5% CO_2_ and pseudoviruses were collected every 24 h and stored at 4°C.

A549/ACE2 cells were seeded in 48-wells plates, cultured for 24 h at 37°C with 5% CO_2_ and transduced with pseudoviruses harboring VSV-G or S-SARS-CoV-2 proteins or lacking the fusion protein (∆Env) in the presence of polybrene (4 µg/ml; Sigma-Aldrich, Poland) and HTCC-77 (100 μg/ml) or control PBS. After 4 h incubation at 37°C unbound virions were removed by washing thrice with 1 × PBS and cells were further cultured for 72 h at 37°C with 5% CO_2._ Cells were lysed in RIPA buffer (50 mM Tris, 150 mM NaCl, 1% Nonidet P-40, 0.5% sodium deoxycholate, 0.1% SDS, pH 7.5) and transferred onto black 96-wells plates. Fluorescence levels were measured on a microplate reader Gemini EM (Molecular Devices, UK).

### Statistical analysis

All the experiments were performed in triplicate, and the results are presented as mean ± standard deviation (SD). To determine the significance of the obtained results Student t test was carried out. P values of < 0.05 were considered significant.

## ACKNOWLEDGEMENTS

This work was supported by the subsidy from the Polish Ministry of Science and Higher Education for the research on the SARS-CoV-2 and a grant from the National Science Center UMO-2017/27/B/NZ6/02488 to KP.

The funders had no role in study design, data collection, and analysis, decision to publish, or preparation of the manuscript.

The technology is owned by the **Jagiellonian University** (Krakow, Poland) and protected by a patent no **WO2013172725A1** and associated documents.

## AUTHOR CONTRIBUTIONS STATEMENT

A.M., Y.C., A.S., E. B. D., X.G., Y.G., J.L., L.C. conducted the experiments. M.O., M.U., and S.R.M provided materials and methods for the study. A.M., K.P. designed the study and experiments, analysed the data and wrote the manuscript. K.L., D.L., F.Z., M.N., K.S. analysed the data. K.P. supervised the study. All authors reviewed the manuscript and approved the submitted version. All authors agreed to be personally accountable for their own contributions and to ensure that questions related the accuracy or integrity of any part of the work are appropriately investigated, resolved, and the resolution documented in the literature.

